# Revealing capture sites and movements by strontium isotope analyses in bones of *Caiman yacare* in the Beni river floodplain, Bolivia

**DOI:** 10.1101/2021.04.14.439857

**Authors:** Marc Pouilly, Sergio Gomez, Christophe Pécheyran, Sylvain Bérail, Gustavo Alvarez, Guido Miranda-Chumacero

## Abstract

Studying the distribution of organisms and their movements is fundamental to understand population dynamics. Most studies indicated that crocodilians do not move around much but several studies demonstrated that some species showed movement patterns. Detection of these movements along the individual life is still a challenge. In this study we analyzed the variation of strontium isotopic ratio (^87^Sr^/86^Sr) in the femur bones of 70 *Caimanya care* individuals caught in 16 sites located in five hydrological sectors of the Beni river floodplain in Bolivia. Our results demonstrated for the first time that such a methodology could yield indications about the capture sites and reconstruct individual life history. Analyses of the outer part of the femur of 70 individuals showed that capture sites could be differentiated between sectors and even between sites or groups of sites in each sector. Studies of complete ^87^Sr^/86^Sr profiles along the femur, representing the individual’s entire life, were performed on 33 yacares. We found that most of the individuals did not show any significant isotopic variation throughout their lives. This absence of variation could result from a high fidelity to the birth site, and/or from an insignificant isotopic variation between the water bodies through which the animal has potentially moved. However, 24% of the analyzed individuals presented significant variations that can be considered as movements between different habitats. Based on the observed low proportion of moving yacares, we advocated that each water body should be considered an individual management unit.

## 1. INTRODUCTION

Studying the distribution of organisms and their movements is fundamental to understand population dynamics (Nathan et al. 2008) and to draw effective conservation and management plans. Although most studies indicate that crocodilians do not migrate much for most life-history stages (Kay 2004), several studies demonstrated that some species showed movement patterns. Indeed, home range size appears strongly influenced by topography, reproductive status, season, and individuals’ length (Brien et al. 2008). Movements could occur on short distance when related to feeding or seasonal water level changes, or on a larger distance when related to reproduction or dispersal, to allow a decrease in demographic density or to avoid predators (hunting pressure for example).

Leslie (1997) and Swanepoel (1999) observed seasonal movements up to 36 km of Nile crocodiles in response to spatial and temporal changes in prey abundance. Tucker et al. (1997) indicate that pubescent males of *Crocodylus johnstoni* were essentially nomadic. Kay (2004) showed movements of *Crocodylus porosus* up to 60 km to nesting habitat for the females and up to 85 km for the males. Kay (2004) also reported that a translocated juvenile male traveled 118 km in 12 days to return to its capture area. In South America, one of the most extensive works on *Caiman yacare* movements reported that the animals moved up to 16 km for females and 18 km for males in a five years period in the vast floodplain of the Brazilian Pantanal (Campos et al. 2006). In Bolivia, De La Quintana et al. (2020) estimated a maximum monthly home range of 19.96 ha for *Melanosuchus niger* and of 1.74 ha for *C. yacare.* These studies remain scarce and are generally limited to a small number of individuals (for example 47 in Campos et al. 2006; 9 for *M. niger* and 3 for *C. yacare* in De La Quintana et al. 2020) and a short period of life because of the methodological and logistical difficulties associated to the use of conventional methods such as capturing-marking-recapturing and telemetry.

Strontium is a trace alkali-earth element that has four naturally-occurring isotopes, among which three are stable (^84^Sr, ^86^Sr and ^88^Sr), and one is radiogenic (^87^Sr) and produced by the radioactive decay of rubidium (^87^Rb, with a half-life of 4.88 x 10^9^ years, Faure & Mensing, 2005). Strontium isotopic composition depends on the nature and age of rocks (Palmer and Edmond, 1992). Generally, carbonate rocks and evaporites rocks present low ^87^Sr/^86^Sr values, whereas silicate minerals are more radiogenic (higher ^87^Sr/^86^Sr). In freshwater systems, ^87^Sr/^86^Sr in the dissolved fraction of water is considered to result from the mixture of Sr from the different rock types present in the drainage area. In the Amazon basin the Sr isotopic differences among various drainage systems are correlated to the rock types in their sediment source regions (Palmer and Edmond, 1992; Gaillardet et al., 1997; Viers et al., 2008, Santos et al. 2015). On a large scale, there is a clear distinction between the isotopic ratios measured in the Madeira River basin, which are more radiogenic (> 0.715), and those measured in the Solimões River basin (< 0.710, Santos et al. 2015). In the Beni river, part of the Madeira watershed, ^87^Sr/^86^Sr differences were also observed on a smaller scale between the mainstream (0.716-0.720), draining Andean waters, and the floodplain lakes (0.725), mostly supplied by local drainage (Pouilly et al. 2014).

Due to its chemical propriety strontium substitutes for calcium in geological and biological matrices. Strontium naturally enters in the composition of tissues of organisms, which incorporates particular elements via water and food. During this incorporation, ^87^Sr/^86^Sr ratio has been shown to remain unchanged from one trophic level to another in the food web, and to reflect the ^87^Sr/^86^Sr of the dissolved fraction of water (e.g. Kennedy et al. 2000 and Pouilly et al. 2014 for Amazonian freshwater fishes). Due to these characteristics, ^87^Sr/^86^Sr could be used as a spatial geochemical marker in freshwater systems presenting geological contrasts.

Many organisms form calcified structures that grow all along with their life. This is the case for example of the bivalve shells, the fish vertebrate bones, scales and otoliths. These structures generally present a continuous growth, so that they are commonly used to determine the age of a specimen by counting the successive growth lines deposited throughout organisms’ life. Microchemical variations can be analyzed at the scale of these growth lines, from the birth to the death of the organisms, to retrieve information about the environment in which the organism lived at each stage of its life. For instance, analyses of ^87^Sr/^86^Sr variation in fish scales or otolith have been used to reconstruct fish movements in the Amazon basin (Sousa et al. 2015, Duponchel et al. 2016, Hauser et al. 2020). It is likely that the same methodology could be applied to evaluate the movement patterns of other aquatic organisms such as crocodilians or river turtles.

The present study aims to determine the capture site and the movement patterns of *Caiman yacare* (Daudin, 1801) in the Beni floodplain, based on strontium isotopic (^87^Sr/^86^Sr) variations along femur bone transects, representing the individual’s life.

We collected yacares in 16 lakes belonging to five hydrological sectors. We draw the hypothesis that 1) the outer borders of yacare’s bones (corresponding to the last period of their life) record the ^87^Sr/^86^Sr values of the lake in which they lived, and that 2) each hydrological sector and site present a specific signature for strontium isotopic ratio. Upon validity, these hypotheses allow using Sr isotopes to determine the capture site of any individual. Moreover, based on an analysis of ^87^Sr/^86^Sr variations along a complete transect on the bones, representing the individuals’ entire life, we tested the hypothesis that yacares present limited movements, and do not move across lakes or sectors during their life.

## Method

### Study area

The Beni River is a major tributary of the Madeira River, in the south west part of the Amazon basin (figure 1). Upper catchments of the Beni River drain the Andean and sub-Andean mountains of northern Bolivia. The corresponding rivers join in the region of Rurrenabaque before entering in a sizeable sedimentary floodplain composed of the Beni River mainstream, many marginal lakes (corresponding to natural tectonic depressions, locally named ‘lago’, or to oxbow lakes, old river beds more or less disconnected from the actual mainstream, named ‘laguna’) and some local tributaries (corresponding to stream, locally named ‘arroyo’).

**FIGURE 1.**
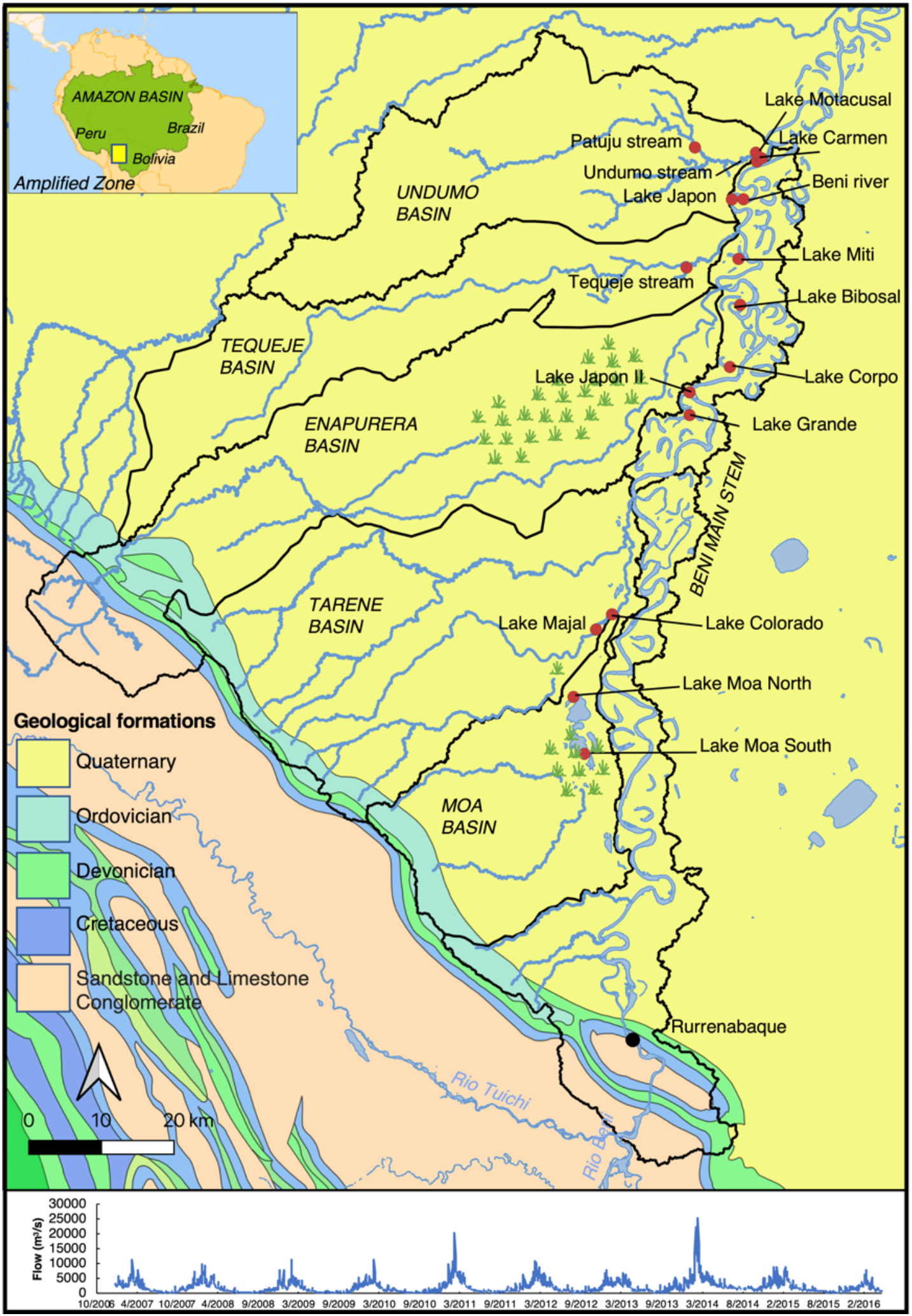
Geological and hydrographic map of the lower subandean and upper lowland areas of the Beni River basin (northern Bolivia). Red circles marked the 16 sites of capture. Green symbol areas correspond to wetlands without a well-defined drainage. Bottom: hydrograph of the Beni river from 2007 to 2017 at the Rurrenabaque gauging station (Data source: www.hybam.org.fr)

The Beni-Madeira basin drains Ordovician and Precambrian rocks of the Andes (Ramos, 2008). Dellinger et al. (2015) measured a high range of variability of ^87^Sr/^86^Sr ratio (0.713-0.733) among the upper tributaries of the Beni river. Santos et al. (2015) observed large seasonal fluctuations in the ^87^Sr/^86^Sr ratio (0.716-0.720) in the Beni River at the Rurrenabaque gauge station, indicating that the sources of dissolved matter coming from the upstream part of the basin vary over time, depending of the relative contribution of each tributary. Pouilly et al. (2014) observed higher ^87^Sr/^86^Sr (0.725) in the marginal floodplain lakes close to the river, and Dellinger et al. (2015) indicate an ^87^Sr/^86^Sr value of 0.726 for one tributary endogen to the floodplain. These contrasts show that while the floodplain is geologically homogeneous, differences in ^87^Sr/^86^Sr of water can be expected depending on the origin of the water. Indeed, in this area, lakes could be supplied by a mix of water from the Beni river mainstream supplied by Andean and sub-Andean drainage areas, and from tributaries originating in the last mountain range before the floodplain or in the local floodplain drainage area (figure 1).

### Yacare capture and sampling

Yacare femur bone samples, that can be used in crocodilian age determination (Hutton 1986, Roberts et al. 1988, Seltzer et al. 2006), were obtained during operations of authorized commercial hunting carried out in 2010, 2016 and 2017 by *Matusha Aidha* Caiman Management Association with the Wildlife Conservation Society support. Operations took place in the Tacana Indigenous Territory divided in five hydrological sectors (figure 1): the active band close to the Beni river (less than 1 km from the Beni main channel) and four tributary basins in the left bank of the Beni River (up to 1.5 km from the Beni River main channel): Moa, Tarene, Tequeje and Undumo basins (from upstream to downstream position, respectively).

After the capture, the total length, weight and sex of the animal were recorded and part of the femur was extracted to estimate age (Gomez 2018). The femurs were kept dry without the meat and transported to the laboratory for preparation. Once in the laboratory, bones were transversally cut with a low-speed saw to obtain a tiny slice of 3-5 mm of thickness. Slices were cut to obtain small bands perpendicular to growth marks (figure 2). Samples were washed, sonicated in distilled water and dried before analyses.

**FIGURE 2.**
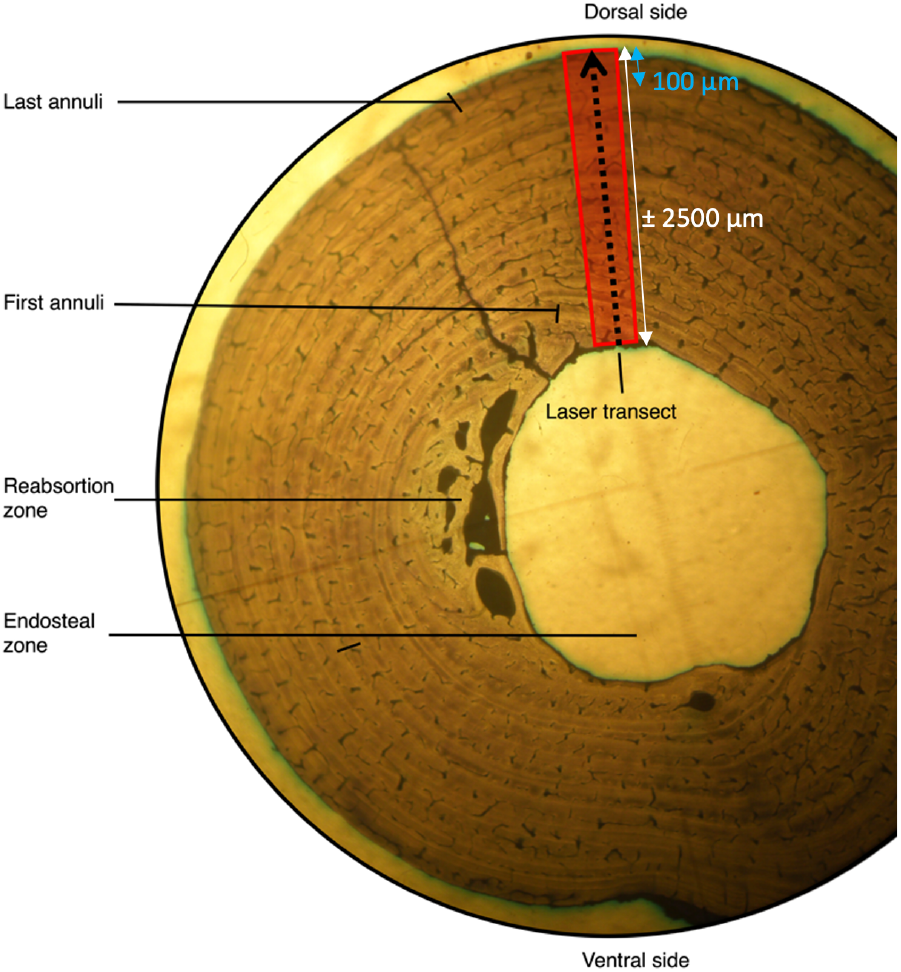
Yacare femur transversal section showing the successive growth lines (annuli) composed by alternation of translucid (low seasonal growth) and obscure (high seasonal growth) layers. Red rectangle and arrow show the internal (birth) – external (death) transect performed by LA-fs-MC-ICPMS to analyse ^87^Sr/^86^Sr variation along the yacare life. White line materialized the full transect analysis, blue line the outer part.

### Sr Isotopic Analyses

Sr isotope analyses were performed at the *Laboratoire de Chimie Analytique Bio-Inorganique et Environnement* (LCABIE) from the *Institut Pluridisciplinaire de Recherche sur l’Environnement et les Matériaux* (IPREM), *Université de Pau et des Pays de l’Adour,* France. The isotope ratios were measured using a Nu Plasma HR-MC-ICPMS (Nu instruments, Wrexham, United Kingdom) coupled with a laser ablation Alfmet femtosecond (fs) (Nexeya SA, Canejan, France) following the procedure detailed by Claverie *et al*. (2009) and Tabouret *et al*. (2010). Laser ablation conditions were 1000 Hz frequency, 20 μJ pulse energy until the depth limit ablation (<30 μm), beam spot size of 20 μm, and velocity 10 μm.s^-1^. The material ablated by the laser was carried with He gas to a double spray chamber in which it was mixed with a 2% HNO3 solution before introduction into the plasma (Barats *et al*. 2007). The interferent ^87^Rb signal was monitored by ^85^Rb, and ^87^Sr/^86^Sr was corrected following Barnett-Johnson *et al.* (2010). Similarly, ^83^Kr was measured to control the impact of ^84^Kr and ^86^Kr on ^84^Sr and ^88^Sr values, respectively. Finally, the ratio ^86^Sr/^88^Sr was used to correct the instrumental mass bias on the ^87^Sr/^86^Sr ratio using the exponential law (Walther and Thorrold, 2008). A reference material (NIESS 22, certificated by National Institute of Japan Environmental Studies) was analyzed at the beginning and at the end of each session to check the measurement accuracy and reproducibility.

In order to address our first objective (to determine the ability of the strontium isotope ratio to indicate the site of capture), analyses were performed on a transect perpendicular to growth marks running on the final external part of the bone (around 100 μm, corresponding to less of one years of adult yacare’s life, figure 2). These analyses resulted in series of values, time related, which records the environmental condition during the last period of the yacare life, before its capture. We used the mean of ^87^Sr^/86^Sr ratios on 100 μm of each sample in Anova tests between yacares from the same site collected in the different years in order to verify that no significant difference could be observed among years. Furthermore, in order to evaluate the geographical scale at which ^87^Sr^/86^Sr could be discriminant, Anova tests were performed between the yacares ^87^Sr/^86^Sr means grouped according to hydrological sectors and between sites of a same sector. All the statistical tests were performed using the R statistical computing freeware program (http://www.r-project.org/). The type I error was set to p-value = 0.05 for all the tests.

In addition, on randomly selected samples, the Sr isotope ratio profile of variation was obtained by a continuous analysis on a transect perpendicular to growth marks running from the bone center (corresponding to the beginning of life) to the bone external border (corresponding to the capture period) (Gomez 2018; Seltzer et al. 2006). These complete transects were carried out to detect ^87^Sr/^86^Sr temporal variations, which could be interpreted as movement through the life of the individual.

## Results

A total of seventy samples were collected in three hunting operations during the dry season: in 47 in 2010, 15 in 2016, and 8 in 2017. Samples proceeded from 16 sites (figure 1, table 1): 7 sites in the active band close to the Beni river, 2 sites in the Moa basin, 2 in the Tarene basin, 1 in the Tequeje basin and 4 sites in the Undumo basin. Sampled yacares were mostly males (68) and ranged in size from 160 to 239 cm in total length, with estimated ages between 7 and >20 years, according to a Von Bertalanffy model calculated by Gomez 2018, (Table S1).

**Table 1.** Total length (min, max), mean (min, max) ^87^Sr/^86^Sr on femur external part and full transect of *Crocodilus yacare* from 16 sites of the lower Beni River floodplain, Bolivia.

| Basin | Total individual | Individual Total Length (cm) | | Final external part $^{87}\text{Sr}/^{86}\text{Sr}$ | | | Full transect $^{87}\text{Sr}/^{86}\text{Sr}$ | | | |
| --- | --- | --- | --- | --- | --- | --- | --- | --- | --- | --- |
|  | number | Min | Max | Mean | Min. | Max. | number | Mean | Min. | Max. |
| **Moa** | **9** | **191** | **223** | **0.7257** | **0.7211** | **0.7285** | **3** | **0.7274** | **0.7244** | **0.7292** |
| Lago Moa Norte | 4 | 191 | 213 | 0.7252 | 0.7232 | 0.7269 | 1 | 0.7244 | 0.7244 | 0.7244 |
| Lago Moa Sur | 5 | 194 | 223 | 0.7260 | 0.7211 | 0.7285 | 2 | 0.7288 | 0.7285 | 0.7292 |
| **Tarene** | **21** | **160** | **237** | **0.7251** | **0.7200** | **0.7266** | **12** | **0.7258** | **0.7243** | **0.7268** |
| Laguna Colorado | 11 | 178 | 237 | 0.7262 | 0.7259 | 0.7266 | 7 | 0.7260 | 0.7252 | 0.7268 |
| Laguna Majal | 10 | 160 | 234 | 0.7240 | 0.7200 | 0.7262 | 5 | 0.7255 | 0.7243 | 0.7260 |
| **Tequeje** | **7** | **174** | **200** | **0.7213** | **0.7203** | **0.7232** | **3** | **0.7213** | **0.7203** | **0.7231** |
| Arroyo Tequeje | 7 | 174 | 200 | 0.7213 | 0.7203 | 0.7232 | 3 | 0.7213 | 0.7203 | 0.7231 |
| **Undumo** | **13** | **187** | **230** | **0.7232** | **0.7218** | **0.7268** | **5** | **0.7233** | **0.7221** | **0.7244** |
| Arroyo Patuju | 3 | 187 | 189 | 0.7239 | 0.7236 | 0.7246 | 3 | 0.7240 | 0.7236 | 0.7244 |
| Arroyo Undumo | 3 | 200 | 229 | 0.7228 | 0.7218 | 0.7236 | 1 | 0.7225 | 0.7225 | 0.7225 |
| Laguna Carmen | 5 | 200 | 230 | 0.7224 | 0.7221 | 0.7227 | 1 | 0.7221 | 0.7221 | 0.7221 |
| Laguna Motacusal | 2 | 194 | 198 | 0.7248 | 0.7228 | 0.7268 |  |  |  |  |
| **Beni** | **20** | **176** | **239** | **0.7218** | **0.7198** | **0.7290** | **9** | **0.7231** | **0.7201** | **0.7302** |
| Laguna Bibosal | 4 | 196 | 222 | 0.7229 | 0.7205 | 0.7290 | 2 | 0.7257 | 0.7213 | 0.7302 |
| Laguna Corpo | 2 | 199 | 202 | 0.7240 | 0.7220 | 0.7261 | 1 | 0.7260 | 0.7260 | 0.7260 |
| Laguna Grande | 3 | 212 | 239 | 0.7203 | 0.7200 | 0.7207 |  |  |  |  |
| Laguna Japon | 1 | 203 | 203 | 0.7217 | 0.7217 | 0.7217 |  |  |  |  |
| Laguna Japon II | 2 | 176 | 194 | 0.7208 | 0.7198 | 0.7219 |  |  |  |  |
| Laguna Miti | 5 | 178 | 200 | 0.7220 | 0.7198 | 0.7232 | 3 | 0.7231 | 0.7229 | 0.7234 |
| Rio Beni | 3 | 182 | 192 | 0.7209 | 0.7203 | 0.7212 | 3 | 0.7204 | 0.7201 | 0.7206 |

### Geographical discrimination by ^87^Sr/^86^Sr

Mean ^87^Sr/^86^Sr ratios of yacare were significantly different between the five hydrological sectors. A clear differences appeared between the two upstream sectors (Moa: 0.7257; Tarene: 0.7251) and the three downstream sectors (Tequeje: 0.7213; Undumo: 0.7232, Beni mainstem: 0.7218; Anova test F=38.6, *p-value* <0.0001, table 1, Figure 3).

**FIGURE 3.**
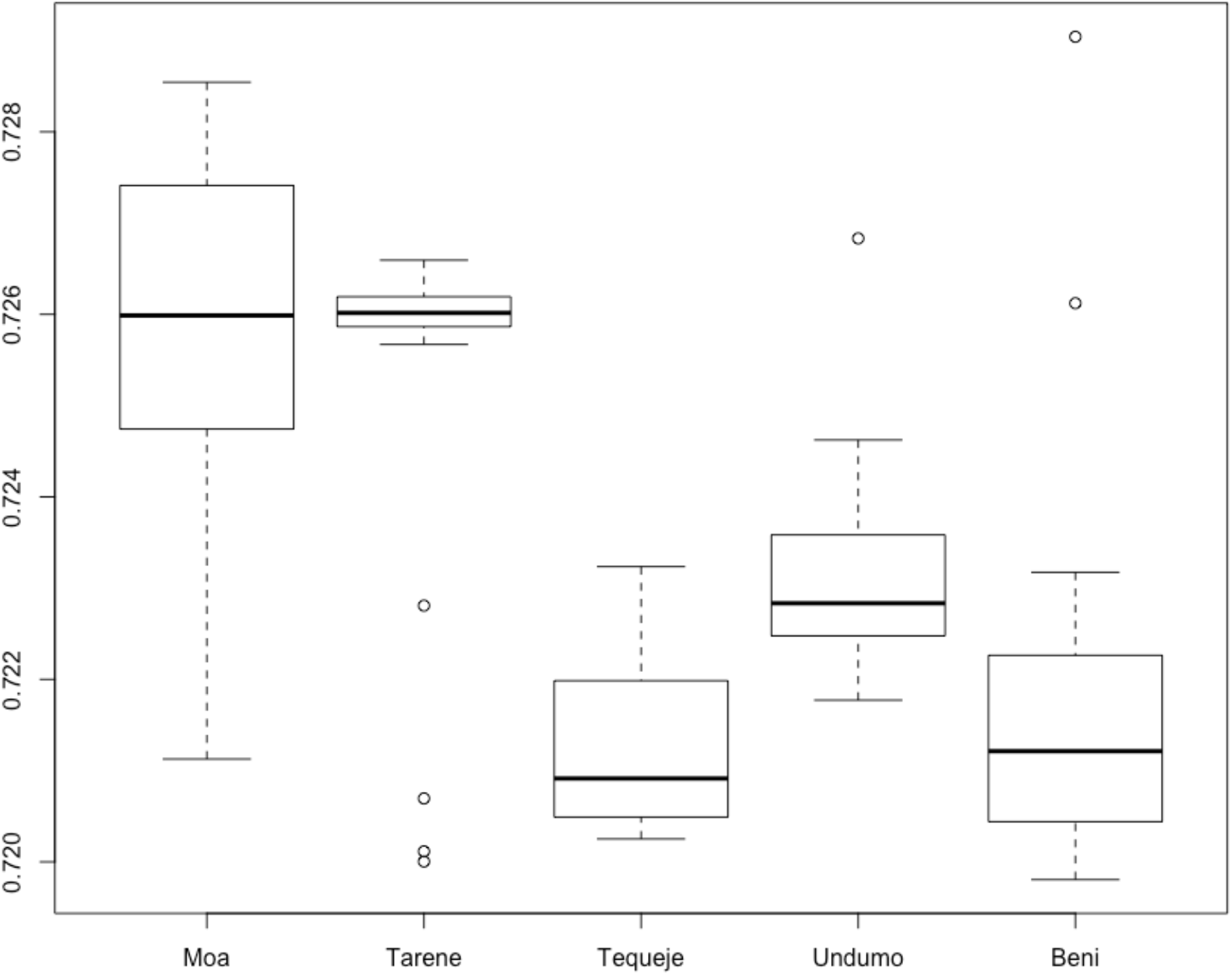
Boxplot of mean ^87^Sr/^86^Sr values of the external part of femurs of yacare captured in five sectors of the lower Beni River floodplain. Yacares presented significantly higher values ^87^Sr/^6^Sr in the Moa and Tarene basins, located upstream in the studied area.

Differences in mean ^87^Sr/^86^Sr were also observed between yacares captured in different sites or group of sites from the same hydrological sector. For example, yacares from the two sites of the Tarene basin presented a significant difference (Laguna Colorado 0,7262 and Laguna Majal 0,7240; F= 7.51, *p-value*=0.013, Figure 4b). In the Undumo basin, yacares presented a significant difference in mean ^87^Sr/^86^Sr (F= 8.20, *p-value*=0.015) between two sites, Arroyo Undumo and Laguna Carmen (0.7228 and 0.7224 respectively), and the two other sites Arroyo Patuju and Laguna Motacusal (0.7239 and 0.7248, respectively, Figure 4c).

**FIGURE 4.**
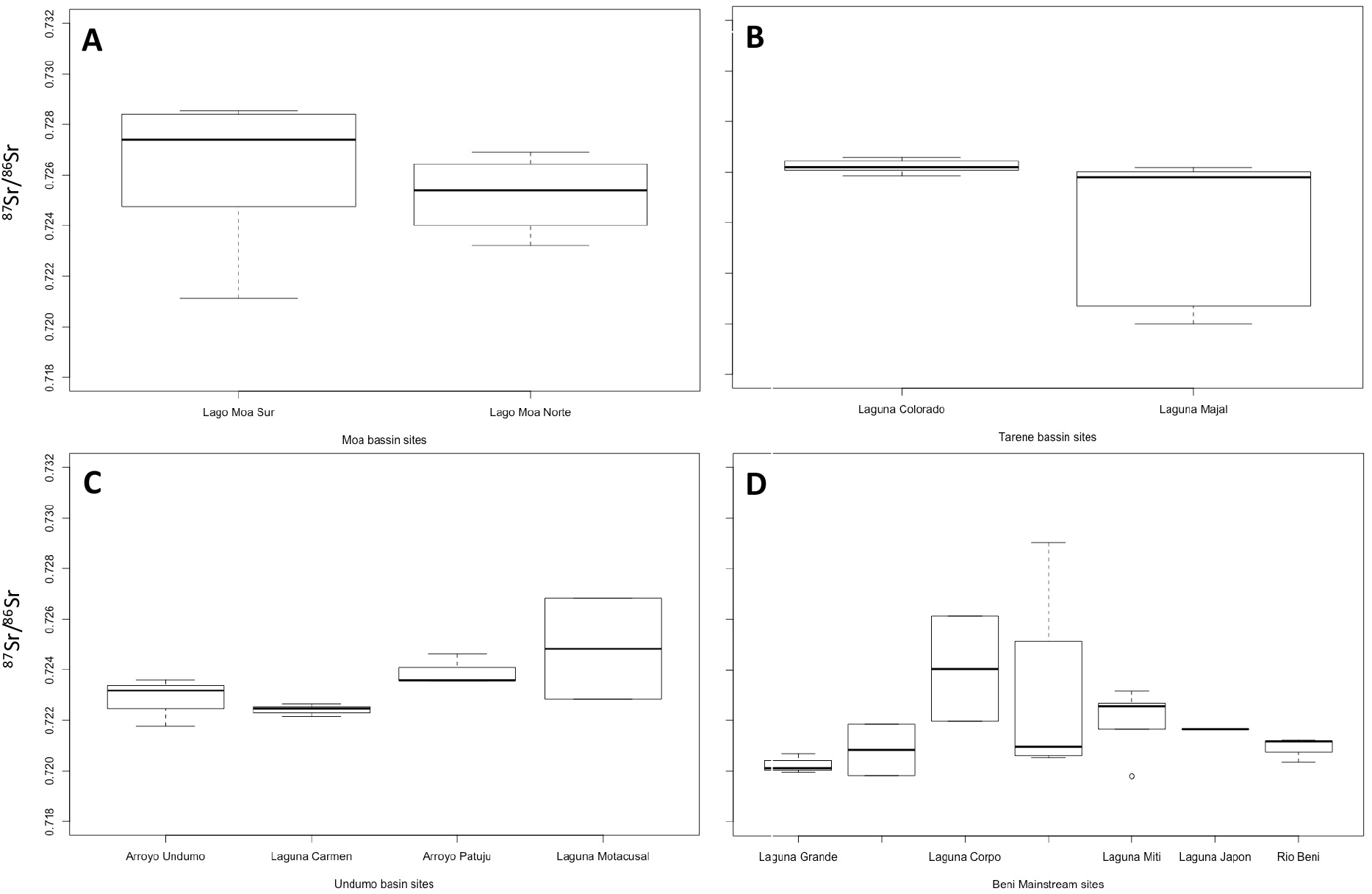
Boxplot of mean ^87^Sr/^86^Sr values of the external part of femurs of yacare captured in different sites inside four watersheds of the lower Beni river floodplain: A- Moa, B- Tarene, C- Undumo, D- Beni. Sites close to the Beni mainstem are ordered according to an upstream-downstream gradient.

Inversely, mean ^87^Sr/^86^Sr of yacares captured in the two sites of the Moa lake, located at the southern and northern ends of the same lake, were not different (0.7260 and 0.7252, respectively; F=0.221, *p-value* > 0.5, Figure 4a). Yacares captured in the active band of the Beni River showed different values depending on the sampling site ranging from 0.7203 (Laguna Grande) to 0.7240 (Laguna Corpo), without clear pattern related to the upstream -downstream gradient or the distance to the mainstream (Figure 4d).

Inside each site, ^87^Sr/^86^Sr presented different degree of interindividual variability (Figure 4). For example, yacares from Laguna Colorado showed low variability (min-max range: 0.7259-0.7266, n=11), whereas those from Laguna Bibosal displayed a higher variability (min-max range 0.7205-0.7290, n=4). Yacares of Laguna Majal individuals presented a bimodal distribution with three yacares characterized by values around 0.7200-0.7206 and six others around 0.7228-0.7262. Similarly, the difference between the southern (Moa Sur, 0.7260) and northern (Moa Norte, 0.7252) parts of the Moa lake is due to the large dispersion of values among the yacares from Lago Moa Sur, with three individuals presenting a mean ^87^Sr/^86^Sr up to 0.7277, and the two others with lower values (0.7247 and 0.7211).

### Movement patterns

Complete profiles of ^87^Sr/^86^Sr were performed from the core to the external border of the femur bone (figure 2) of 33 yacares. Twenty-five presented small variations with no clear trend and that were not significantly outside the range of variability of the sector or the site. The eight other (24%) showed significative variations in the ^87^Sr/^86^Sr ratio along the profile (figure 5): M14 (TL 222cm, 12 years) from Laguna Colorado (Tarene bassin) presented one clear and sharp peak from 0.7260 to 0.7300. M52 (TL 224cm, 14 years) from the same lake presented a clear decrease of ^87^Sr/^86^Sr ratio after 1/3 of the sample from 0.727 to 0.725 before to recover intermediate values (around 0.726). From the same lake, the profile of an individual (4723, TL 237cm, >20 years) also began by low values (0.725) until the middle of the life and increased smoothly to values similar to other individuals from the same lake (0.727). Another Individual (4251, TL 160cm, 7 years) from Laguna Majal in the same Tarene sector, showed a similar profile with low values (0.723) until the middle of the life, but an abrupt shift to higher values similar to those of the other individuals of the lake (0.725). M10 (TL 211cm, 15 years) of Laguna Bibosal, near the Beni mainstem, presented the most outstanding profile among the analyzed individuals. The ^87^Sr/^86^Sr values reach up to 0.730, and the whole pattern is significantly different from all other individuals from the same site. M6 (TL 219cm, 11 years) from Lago Moa Sur showed a continuous decrease from values > 0.730 to 0.727. Finally, two individuals from the Beni River, showed variation from low (0.720) to higher values (0.723). M54 (TL 182cm, 8 years) presents an abrupt change in strontium ratio at the end of the profile, and M48 (TL 183cm, 8 years) displays two peaks also at the end of the transect.

**FIGURE 5.**
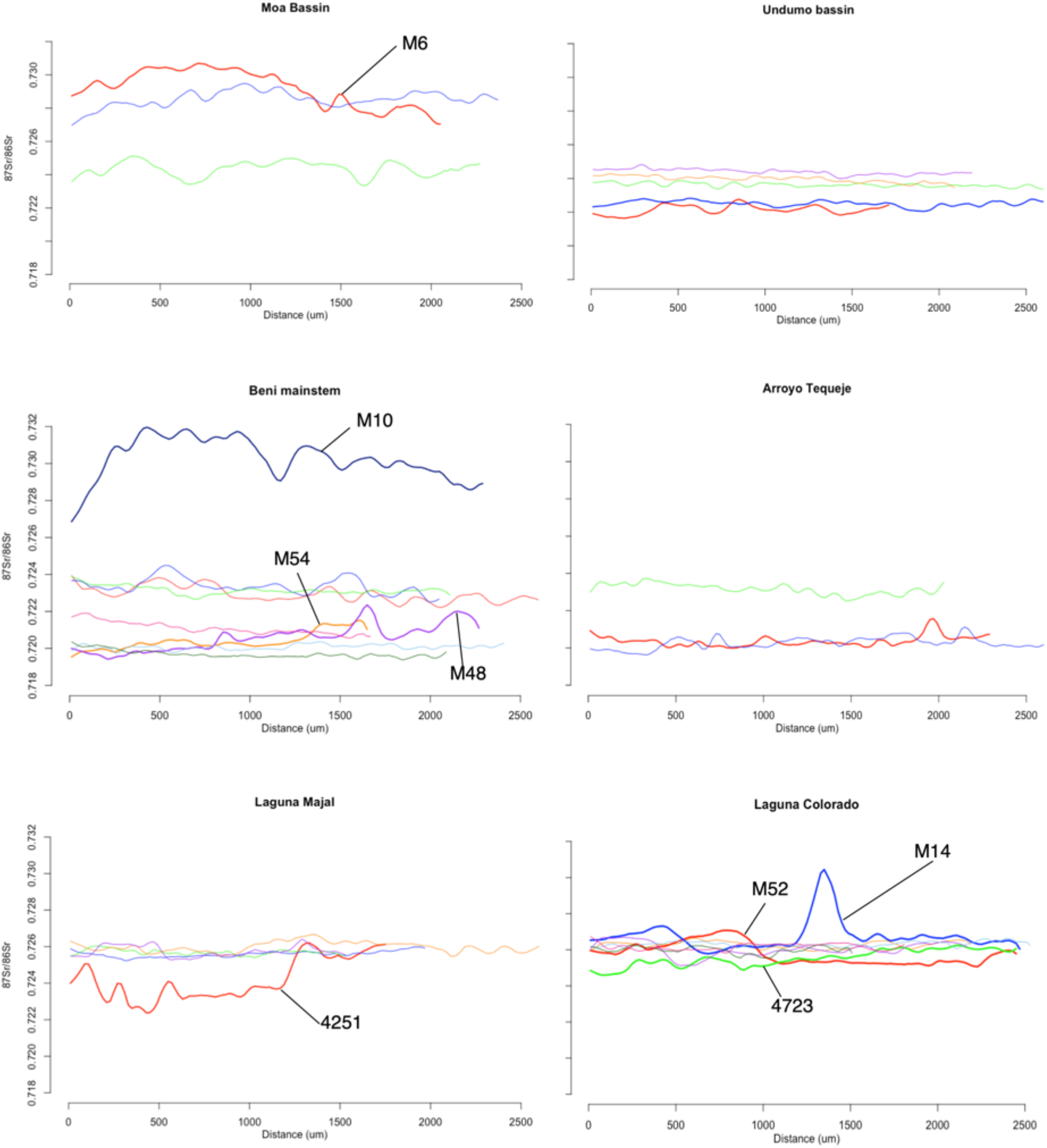
^87^Sr/^86^Sr variation in femur bones (internal - external profiles, representing the individual life history) for 33 yacares captured in several sites of the lower Beni River floodplain. Abscise axis represent the distance to the internal border of the femur (life beginning). Individuals presenting a high variation during their life are identified.

## Discussion

Isotopes, as the ^87^Sr/^86^Sr used in our study, are particularly useful for studying animal movements because they do not require marking or recapturing individuals and they provide time-integrated information that can be linked to geographical regions (Rubenstein and Hobson 2004). However, their effectiveness is primarily linked to the existence of geographic difference of environmental ^87^Sr/^86^Sr ratio due to regional differences in the geologic substrate. Difference of ^87^Sr/^86^Sr in animals could also be attributed to diet (Wheatley et al. 2012). *Caiman yacare* is mainly piscivore (Santos et al. 1996), and fish ^87^Sr/^86^Sr has been shown to be directly related to the environmental isotope ratio (Pouilly et al. 2014). Therefore, it is likely that the ^87^Sr/^86^Sr of *Caiman yacare* bone should be correlated to their environment without a discordant effect of diet. This postulate is widely used in archeological studies (e.g. Bentley 2006) and can be also assume for this study.

Our results demonstrate for the first time that such a methodology could give indications about the localization of capture sites as well as for reconstructing the life history of *Caiman yacare* based on the microchemical analysis of their bones. Moreover, although more work will be necessary to ascertain this finding, the environmental conditions of the Beni river floodplain appeared conducive enough to allow for a geographic discrimination at hydrological sector’s scale and even, with more caution, in some case at the scale of site or goup of site. As crocodiles in general (Kay 2004) and *Caiman yacare* in particular (De La Quintana et al. 2020, Campos et al. 2006) presented relatively small home ranges, the possibility of movement’s interpretation between sites would be profitable. However, this possibility should be improved by a more detail study.

The Beni River main channel presented a marked seasonal variation in ^87^Sr/^86^Sr ratios (min-max around 0.716-0.720), correlated with water discharge (Santos et al. 2015) as well as a lateral variation in the floodplain (up to 0.727, Pouilly et al. 2014, Dellinger et al. 2015) likely linked to local source of water. Pereira et al. (2019) also observed a high variability of Sr isotopes in the fish species *Arapaima gigas* of the lower Beni River and Madre de Dios Rivers (0.715 - 0.730). These observations indicated that the basin is not homogeneous and has not well mixed sources of Sr. However, the water ^87^Sr/^86^Sr baseline was not available at the hydrological sector or lake level. This lack of information limited the possibility of interpretation of our results which was only based on yacare bones. Nevertheless, assuming that Sr incorporated into bones reflects the bioavailable Sr present in surrounding waters (Kennedy et al. 2002, Koch 2007, Wheatley et al. 2012), it is likely that the ^87^Sr/^86^Sr values obtained in the outermost part of the bones corresponded to the ^87^Sr/^86^Sr values characterizing the direct environment of the yacare during its last period of life. All the studied specimens showed a stable and homogeneous profile of ^87^Sr/^86^Sr ratios before the capture, indicating that they likely remained sedentary during this period (or moved inside habitats with the same range of ^87^Sr/^86^Sr values). These differences in individual values could be interpreted as differences between the sites of capture. The ^87^Sr/^86^Sr differences observed between the yacares from the upstream sector (Moa and Tarene watersheds) and the downstream sector (Tequeje and Undumo watersheds), separated by around 80 km (in straight line, and up to 200 km along the river course), indicated a possible geographic discrimination at this scale. Moreover, the ^87^Sr/^86^Sr differences observed between the yacares from sites of the same sector indicated that the discrimination from ^87^Sr/^86^Sr ratios could also be operative at the scale of a site or a group of sites.

Generally, biological samples are compared to environmental baseline obtained from water or another abiotic component. In cases where such environmental baseline is lacking, such as for the present study, we assume that the external part of yacare femurs could be used as a proxy of the ^87^Sr/^86^Sr environmental baseline itself. Although in our study, all sampling sites are localized in the same geologically homogeneous floodplain, we observed significant difference of ^87^Sr/^86^Sr in external part of yacare values among hydrological sectors and even among sites inside a hydrological sector. These differences could be attributed to different origins of waters (Andean, sub-Andean, local drainage) that may likely generate the ^87^Sr/^86^Sr variabilities in dissolved water necessary to revealed yacares’ movements. Increasing the number of samples proceeding from more sites, would permit to draw a more precise baseline of ^87^Sr/^86^Sr between lakes and then to improve the possibility of movements tracking. Such development would require a test of correspondence between the ^87^Sr/^86^Sr values of waters and yacare bones of the same locations.

Even with a more precise baseline, overlapping of ^87^Sr/^86^Sr values between some sampling sites will probably occurred. Examination of other isotopic proxies like oxygen and sulphur isotope (Goedert et al. 2020) could also be appropriate to resolve the situations of overlap. Improve this geographic information would be of importance for population dynamics analysis, as it makes it possible to determine more precisely if the yacares captured at one site or sector were born in this site or sector or if they moved from one site/sector to another during their life. This information could also be useful for harvesting monitoring as it can served as a proof of the region of capture. These tools of commercial traceability are now growing in importance (Baffi & Trincherini 2016, Pereira et al. 2019). Our results should be improved by a more precise baseline and eventually completed with other biogeochemical tags in order to reach a better level of accuracy.

However, the differences among hydrological sectors and sites are sufficient to explore potential yacares movements. Based on the analysis of complete transects on femur bone ranging from birth to capture period of the yacares, we found that most of the individuals do not show significant ^87^Sr/^86^Sr variation throughout their lives. This absence of variation could be the result of (1) a high fidelity of the individual to its birth site, and/or of (2) an insignificant isotopic variation between the water bodies through which the animal has potentially moved. As we detected significant variation between yacares captured in different sites close to each another, hypothesis (1) above is the most likely. These results supported the observations of Campos et al. (2006) and De La Quintana et al. (2020) that estimated, by telemetry, that yacare’s home range are very limited (between 0.48 ha to 248 ha for a short period of their life).

Topography, habitat diversity and connectivity (lake, wetland, secondary stream and main river), and hydrologic seasonality in the Beni River are conducive to yacare’s movements. In opposition a site fidelity hypothesis, seasonal (short) movements, following the water terrestrial transition zone along the lateral dimension of the river, could be expected, as they allow for finding better reproductive or diet condition. Longer-range movements could also be beneficial as they improve the dispersal of the species. Our results did not support these alternative hypotheses as most of the yacares presented a relative stability throughout their life although significant difference of ^87^Sr/^86^Sr in water existed among hydrological sectors. An illustrative example comes from the Lake Moa which is 8.8 km long and around 2600 ha in area, making it a large lake for the Beni floodplain. Yacares captured in the northern part of the lake have a different isotopic ratio than those from the southern part (mean ^87^Sr/^86^Sr respectively 0.725 and 0.728) and presented stable and non-overlapping profiles. In this particular situation, which may correspond to a lake with different water origins, we can deduce that the home range of a specific individual does not encompass all the lake area and, hence, that yacare does not move at this scale. Home range appears therefore not to be related to the area of the lake, nor to connectivity among other habitats.

The previous affirmations are based on the pattern observed on the majority of the yacares analyzed and we can conclude that most of the studied population presents a sedentary pattern. Femur transversal sections showed an alternance of clear and obscure bands, that likely correspond to seasonal marks related to growth rate differences. Clear bands are generally characteristics of a faster growth and inversely obscure bands correspond to a slower growth during the high flow period. In our samples, thickness of these bands was generally higher (50 to 100 μm) to the laser spot size (20 μm), allowing to detect eventual seasonal variation. None of the analyzed yacares showed a repetitive variation that could be interpreted as a seasonal change. As the main channel of the Beni River shows a high seasonal variability in ^87^Sr/^86^Sr values (Santos et al. 2015) we suggest that yacare colonized preferentially marginal habitats with more stable ^87^Sr/^86^Sr conditions. Yacares captured near the Beni mainstem, and in particular three individuals caught in the Beni River site (an oxbow lake close to the main stream), also present a stable profile with ^87^Sr/^86^Sr values near or up 0.720 that correspond to the seasonal maximum of ^87^Sr/^86^Sr values in the water of the main channel (occurring during the high-water season, Santos et al. 2015). No ^87^Sr/^86^Sr values close to 0.716, corresponding to the minimum value of main channel water during the low-water season was observed in yacares, suggesting that these yacares didn’t moved between the river and the adjacent lake. This result, based only in few individuals, is not concordant with the observations of Campos et al. 2006, who concluded to frequent movements from lake to river. Analyze of yacare’s captured in the river would help to evaluate the proportion of individuals moving between the two habitats.

However, 24% (8 individuals) of the analyzed yacares showed significant variation in their ^87^Sr/^86^Sr profiles. One individual (M14) presented a net temporal change before recovering ^87^Sr/^86^Sr values similar to those the beginning of its life. The other individuals presented a more or less two-step profile (figure 5). Interpretation of these patterns are speculative in the absence of a precise baseline. However, we can suggest that the two-step profiles may correspond to a unique migration from one site to another and that these individuals participate to the dispersion and gene mixing of the population.

In crocodilian populations, larger males are considered as dominant for territory and are thus expected to be more sedentary (Drews 1990). On the contrary since females are considered less territorial, it can be hypothesized that females moved more than males (Campos et al. 2006, De la Quintana et al. 2020). In the Tuichi area, in the subandean part of the Beni basin, a female has been observed moving with all her brood (C. Nay, Personal Observation, Figure 6). Our dataset features only two female individuals, both presenting a stable profile. More females are needed to draw conclusions on the sex ratio of non-sedentary individuals and to interpret the driver of these movements.

**FIGURE 6.**
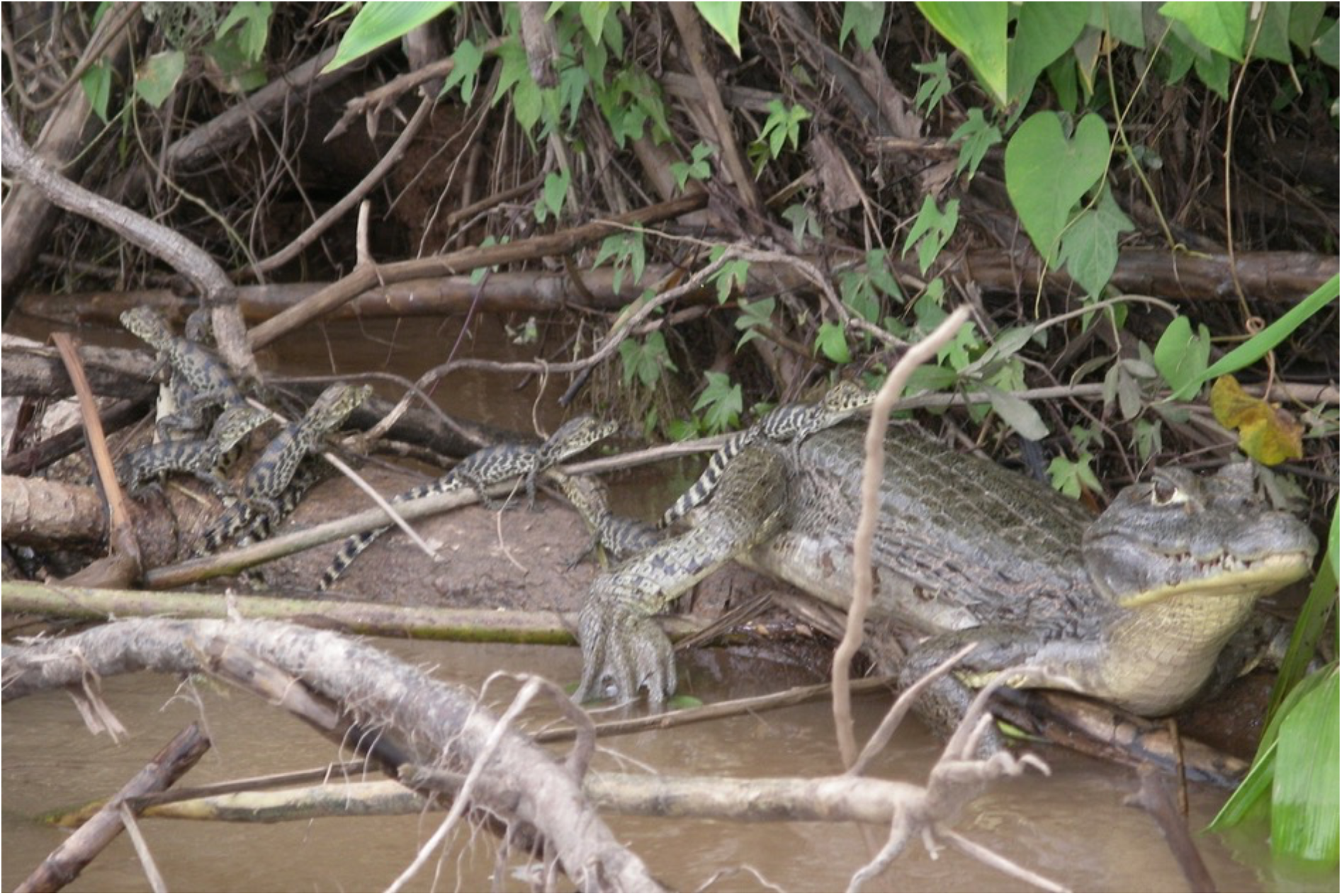
Female yacare on the move with their hatchlings in the Tuichi River, near the study area. Photograph: Constantino Nay (WCS Bolivia).

Campos et al. (2006) suggest that, despite the generally-observed absence of large movement, yacare can travel up to 18 km and frequently move from lake to the river area. They then concluded that management plans should not consider lake and river as autonomous units. Our results indicate that a relative low proportion (<25%) of yacares move. Without denying the need to maintain the possibilities of movement between the different habitats, we suggest that each of them should be considered as an individual autonomous management unit in order to protect the most of the population remaining in the habitat through quotas and minimum catch sizes. Indeed, with small home range and a low proportion of yacares on the move, the probability of restocking an over-hunted site could be low and critical. The conservation of connectivity between habitats should however be targeted in the management plans in order to preserve the dispersal and mixing capacity of the population, considering in particular the movements of females.

## Contributions

MP and GMC designed the study, made the statistical analysis, interpreted the result and redacted the manuscript. GMC, SG and GA made the field sampling and prepare the bones samples. MP, CP and SB performed the strontium isotope analyses. SG, GA, CP, and SB contributed to results preparation and interpretation, and to writing.

## Acknowledgments

We thank the Tacana Caiman Association Matusha Aidha for their help in this study. To Consejo Indigena del Pueblo Tacana (CIPTA) for allowing the work. To Ninon Rios, Magaly Manedoza, Sandra Ribera, Agustin Estivariz, Pablo Justiniano for their help in collecting samples of bones in the caiman harvests. To Julio Pinto and Ruben Marin from the Unidad de Limnologia from UMSA for their help in their laboratory of schlerochronology. To Jesus Nuñez for their help to the laboratory of schlerochronology. To Luis Pacheco (Universidad Mayor de San Andres, Instituto de Ecología, La Paz, Bolivia), Julien Bouchez (Institut de Physique du Globe de Paris, France) and Roberto Santos (Laboratório de Geocronologia, Universidade de Brasilia, Brasil) for the revision and comments on the manuscript.

## Supplementary material

**Table S1:**
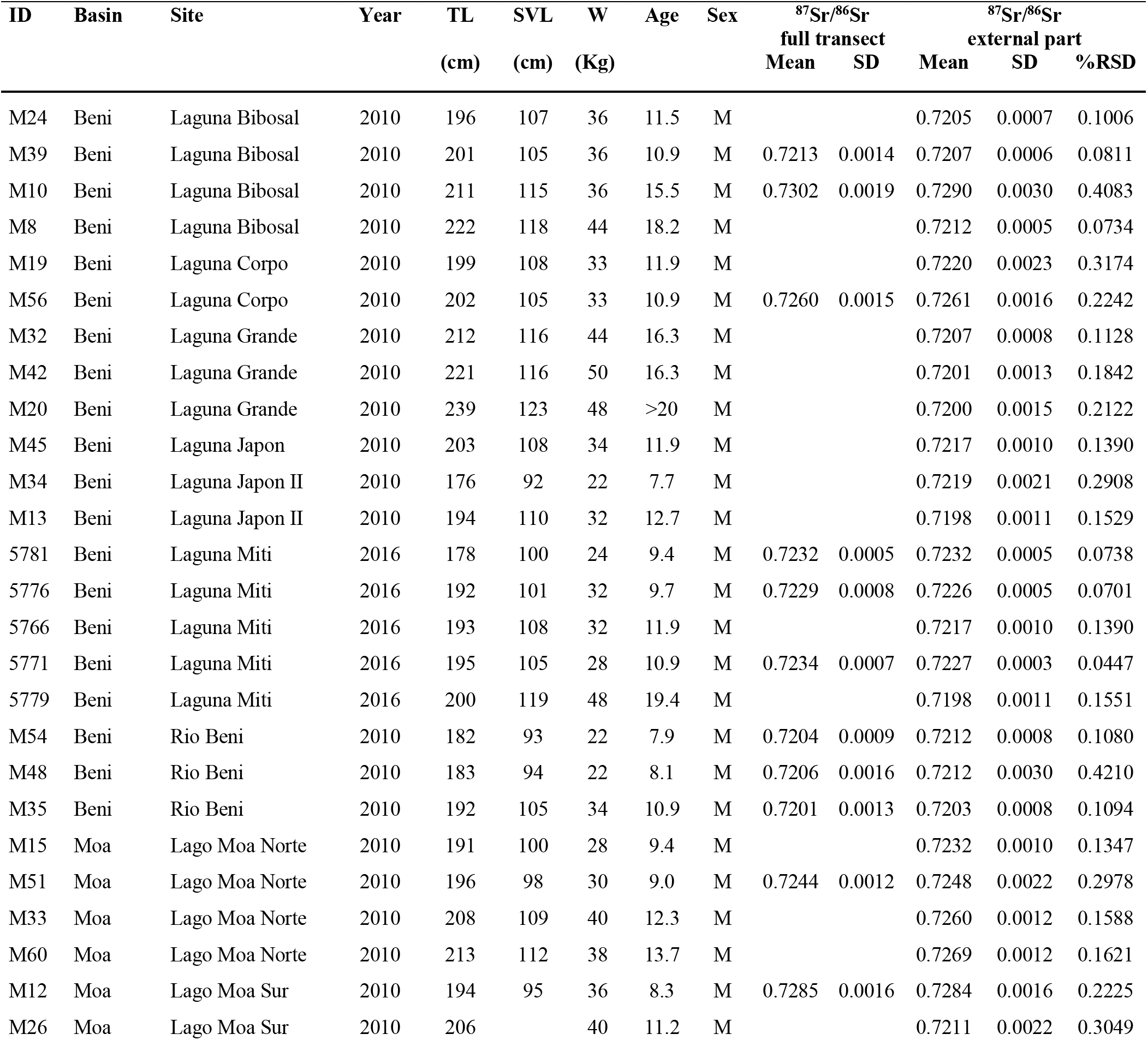
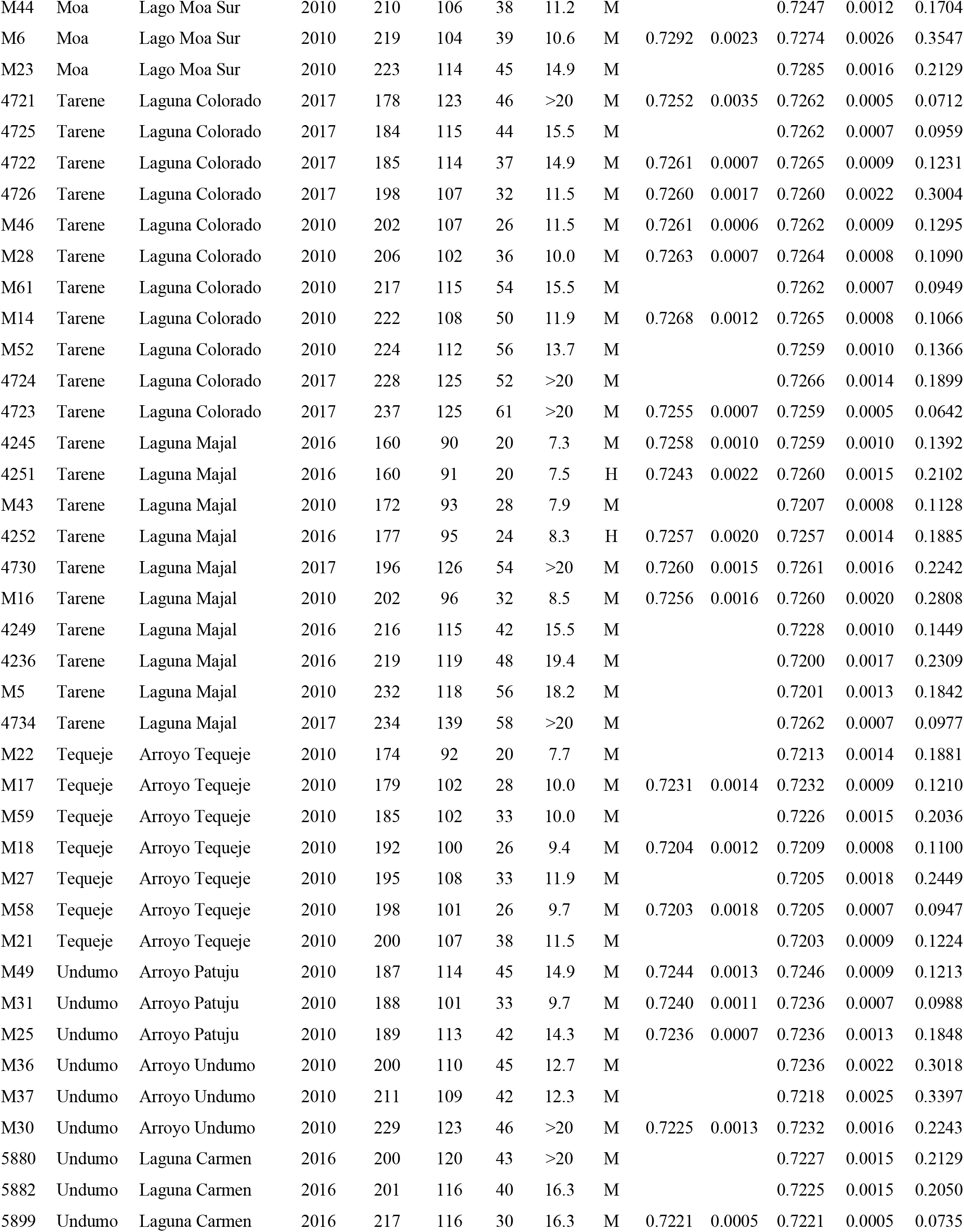
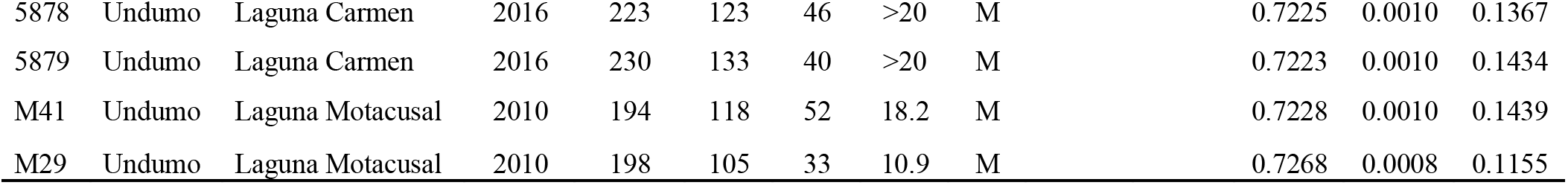
Site, year of capture and individual values of Total Length (TL), Snout Ventral Length (SVL), Weight (W), Age, Sex and ^87^Sr/^86^Sr mean and SD on full transect and external part (100 μm) femur bone. Age column corresponds to estimated values according to a Von Bertalanffy model developed on yacares of the same area by Gomez (2018).

